# Endogenous Plasmids and Reductive Genome Evolution in Host-Associated Bacteria

**DOI:** 10.1101/2022.09.09.507235

**Authors:** Qing Xiong, Cathy Sin-Hang Fung, Xiaojun Xiao, Angel Tsz-Yau Wan, Mingqiang Wang, Yaning Ren, Kevin Yi Yang, Yubao Cui, Xiaoyu Liu, Stephen Kwok-Wing Tsui

## Abstract

Reductive genome evolution is commonly observed among host-associated bacteria including many important pathogens, such as *Mycobacterium leprae* but its molecular mechanism is not well understood ^1–5^. One of the most widely accepted hypotheses to explain bacterial genome reduction is Muller’s ratchet, in which the associated bacteria tend to accumulate deleterious mutations for reduction in the absence of chromosomal recombination inside the eukaryotic host organism ^1,2^. *Cardinium* species belong to the family Amoebophilaceae of the CFB group bacteria, which are a group of endosymbiont bacteria widely distributed among arthropods, that along with *Wolbachia* can cause cytoplasmic incompatibility ^6,7^. In this study, we explored bacterial reductive evolution within the *de novo* assembled genomes of *Cardinium* endosymbionts in two astigmatic mites ^8,9^. Our results shed light on the reduction mechanism driven by endogenous plasmids and their encoded enzymes.

## Main

Using high-throughput genomic sequencing data (Table S1), we *de novo* assembled the genomes of *Cardinium* endosymbionts (Table 1) in two astigmatic mites, *Dermatophagoides farinae* and *Tyrophagus putrescentiae*. The *Cardinium* endosymbiont of *Dermatophagoides farinae, Cardinium sp. DF*, has a genome size of 1,259,597 bp in a single contig (Table 1, Fig. 1A). The genome assembly and the annotated 1,198 proteins were assessed as 76.4% and 77.7% complete, respectively (Table 1). A previous genome assembly of the *Cardinium* endosymbiont of *Dermatophagoides farinae* was constructed in five contigs, but only the longest contig was considered as the genome ^10^ (Table 1). To differentiate the two *Cardinium* endosymbionts of *Dermatophagoides farinae*, the previous assembly was named *Cardinium sp. DF UM* because it was reported by the University of Michigan. The shared identity of the 16s rRNA sequences of the two *Cardinium* endosymbionts of *Dermatophagoides farinae* was 100%. The dot plot of two *Cardinium sp. DF* genome assemblies suggested that both were circular DNA. The assembly of *Cardinium sp. DF UM* was missing the 675,805-676,487 bp region in our assembly (Fig. S1A). Intriguingly, the genome assembly of *Cardinium sp. DF* was annotated with fewer protein-coding genes but higher completeness (Table 1). We confirmed that the low-quality assembly, especially in repetitive bases, caused false-positive frameshifts and gene fission in the genome of *Cardinium sp. DF UM*, which resulted in more genes but lower completeness. Since both genomes were assembled by third-generation sequencing (TGS) long reads ^10^, we proposed that the quality discrepancy was a result of different levels of coverage, although the raw sequencing data of *Cardinium sp. DF UM* are not available.

**Fig. 1.**
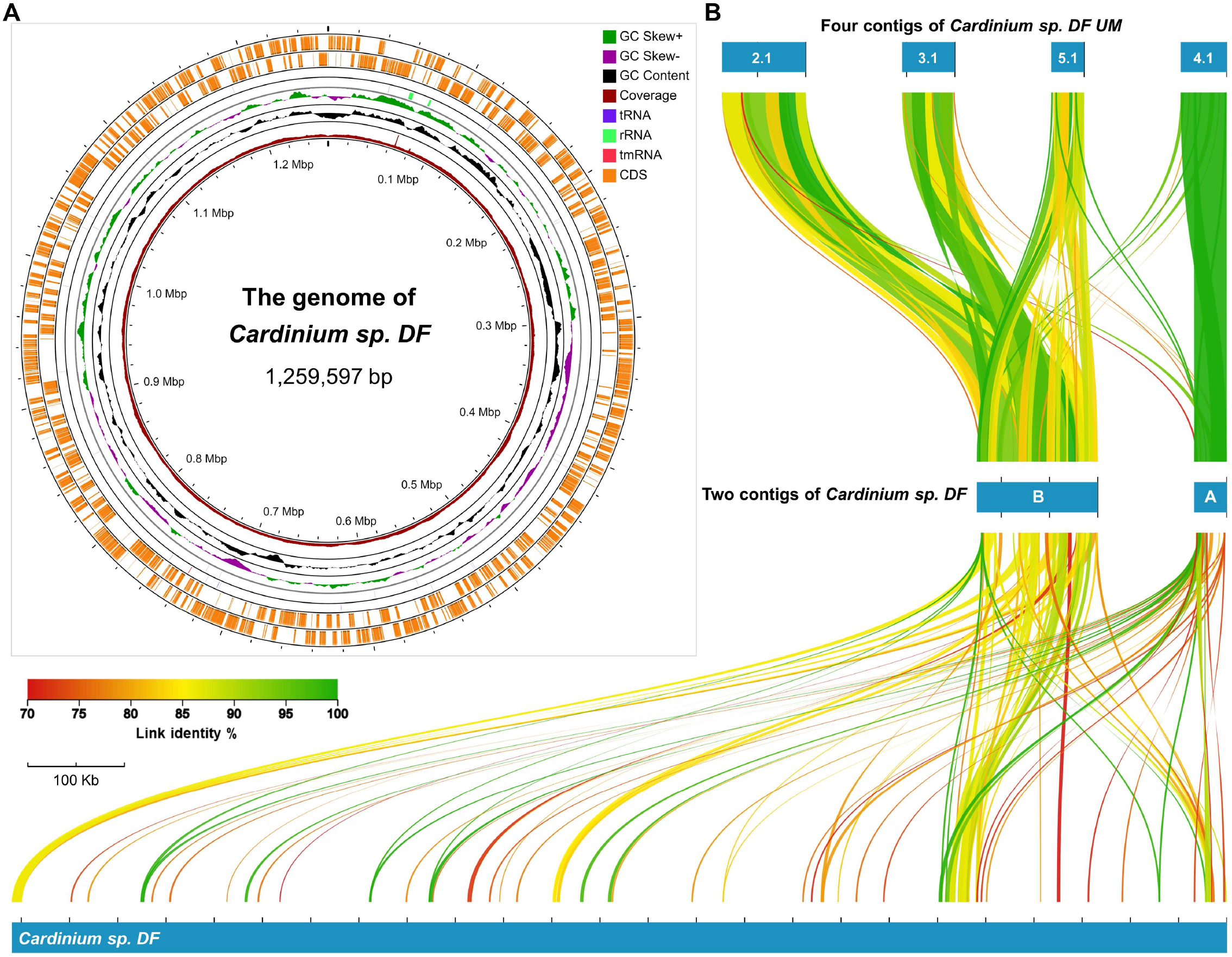
The genome map and whole alignment of *Cardinium sp. DF*. **(A)** Circular genome map of *Cardinium sp. DF*. The genome annotation was performed by Prokka and visualized by the online tool Proksee. The coverage was calculated with the Nanopore and PacBio long reads mapping, in which the height of the most internal ring was computed as (the site coverage)/(the highest coverage) and the full height indicated the highest coverage of 556 X. **(B)** Whole alignment of *Cardinium sp. DF* genome and other short contigs performed by AliTV and further filtered by 70% identity and 2-kb link length. Two short contigs of *Cardinium sp. DF* and four short contigs of *Cardinium sp. DF UM* were aligned with the genome sequence of *Cardinium sp. DF*. Two contigs of *Cardinium sp. DF* are named as contigs A and B. All the contigs of *Cardinium sp. DF UM* have a prefix of NZ_VMBH0100000 in the NCBI GenBank database.

**Table 1.**
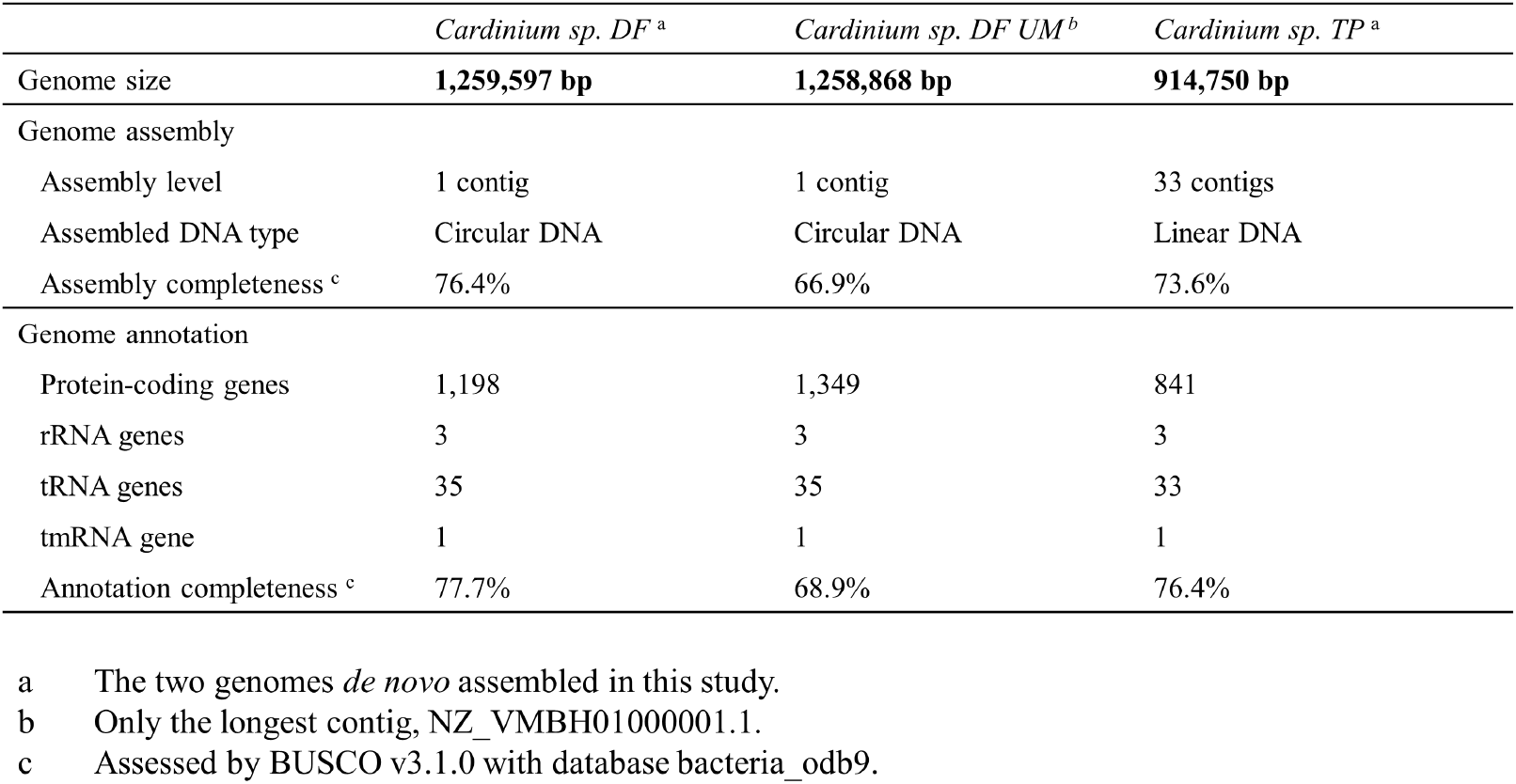
Overview statistics of assembly and annotation of three *Cardinium* bacteria.

For the *Cardinium* endosymbiont of *Tyrophagus putrescentiae, Cardinium sp. TP*, the genome was assembled into 914,750 bp and 33 contigs (Table 1). The completeness of the genome assembly and annotation was 73.6% and 76.4%, respectively. The 16s rRNA of *Cardinium sp. TP* was reported, but there is no available genome assembly ^11^. The low completeness (< 80%) indicated genome reduction during its evolution. The assembly quality of *Cardinium sp. DF* was apparently much better than that of *Cardinium sp. TP* because of the higher sequencing coverage (108.6 X TGS only reads of *Cardinium sp. DF*, <8 X NGS reads of *Cardinium sp. TP* and even fewer TGS reads). Considering their unequal assembly qualities, we cannot conclude that *Cardinium sp. DF* possesses a larger genome size than *Cardinium sp. TP*.

Along with the single contig of *Cardinium sp. DF* (Fig. 1A), two short contigs were assembled and presumed to be extrachromosomal genetic elements (Fig. 1B). Similarly, four short contigs were assembled within the genome of *Cardinium sp. DF UM* (Fig. 1B). In the entire alignment (Fig. 1B), the short contig A was mapped to the contig, NZ_VMBH01000004.1 of *Cardinium sp. DF UM* in high quality, while the other contig B was variably mapped to the other three contigs of *Cardinium sp. DF UM*. Both contigs A and B were dispersed across the genome, as shown in a wide range of highly conserved alignments (cut-off was set as 70% identity and 2-kb length, Fig. 1B), which supports that they are extrachromosomal genetic elements of *Cardinium sp. DF*.

Phylogenetic analysis was performed for the two *de novo* assembled genomes of *Cardinium* endosymbionts, along with other published sequences (Table S2). To generate a high-quality phylogenetic analysis, 10 genome assemblies of *Cardinium* and *Amoebophilus asiaticus* as outgroup were collected for their fewer than 50 assembled sequences (Table S2), annotated, and extracted with 295 single-copy orthogroups (OGs). However, *Cardinium sp. TP* was unexpectedly clustered with the *Cardinium* endosymbiont of *Sogatella furcifera, Cardinium sp. Sogatella furcifera* (Fig. 2A and S2A), in which their protein identity and similarity were 95.62% and 96.58% respectively (Fig. S3). Among nine *Cardinium* genomes, two from *Cardinium hertigii* were located as outgroups to seven other *Cardinium* strains without official species names (Fig. 2A and S2A, Table S2). The other phylogenetic tree based on 16s rRNA sequence was constructed with four additional sequences from oribatid mites, the sister group of astigmatic mites (Fig. S2B). Although the two *Cardinium sp. DF* were clustered with three from oribatid mites, the extremely long branches of the two from *Microzetorchestes emeryi* impeded further discussion (Fig. S2B). Similar to the phylogenetic tree based on 295 single-copy OGs, *Cardinium sp. TP* clustered with *Cardinium sp. Sogatella furcifera* (Fig. S2B). The similarity among *Cardinium* genomes was further explored in whole genome alignment performed by AliTV ^12^ (Fig. S4). The genomic sequences two *Cardinium sp. DF* were mapped and aligned with high quality (Fig. S4). The fragmented genome of *Cardinium sp. TP* could be mapped to that of *Cardinium sp. Sogatella furcifera* with high quality (Fig. S4). *Sogatella furcifera* is well known as an important pest species in rice, while the host of *Cardinium sp. TP*, *Tyrophagus putrescentiae*, is a storage mite mainly infesting stored grains, including rice. The closely related living environments of their hosts could explain why *Cardinium sp. TP* shares a close phylogenetic relationship with *Cardinium sp. Sogatella furcifera* ^13^, but not *Cardinium sp. DF* from the closely related host, house dust mite *Dermatophagoides farinae*.

**Fig. 2.**
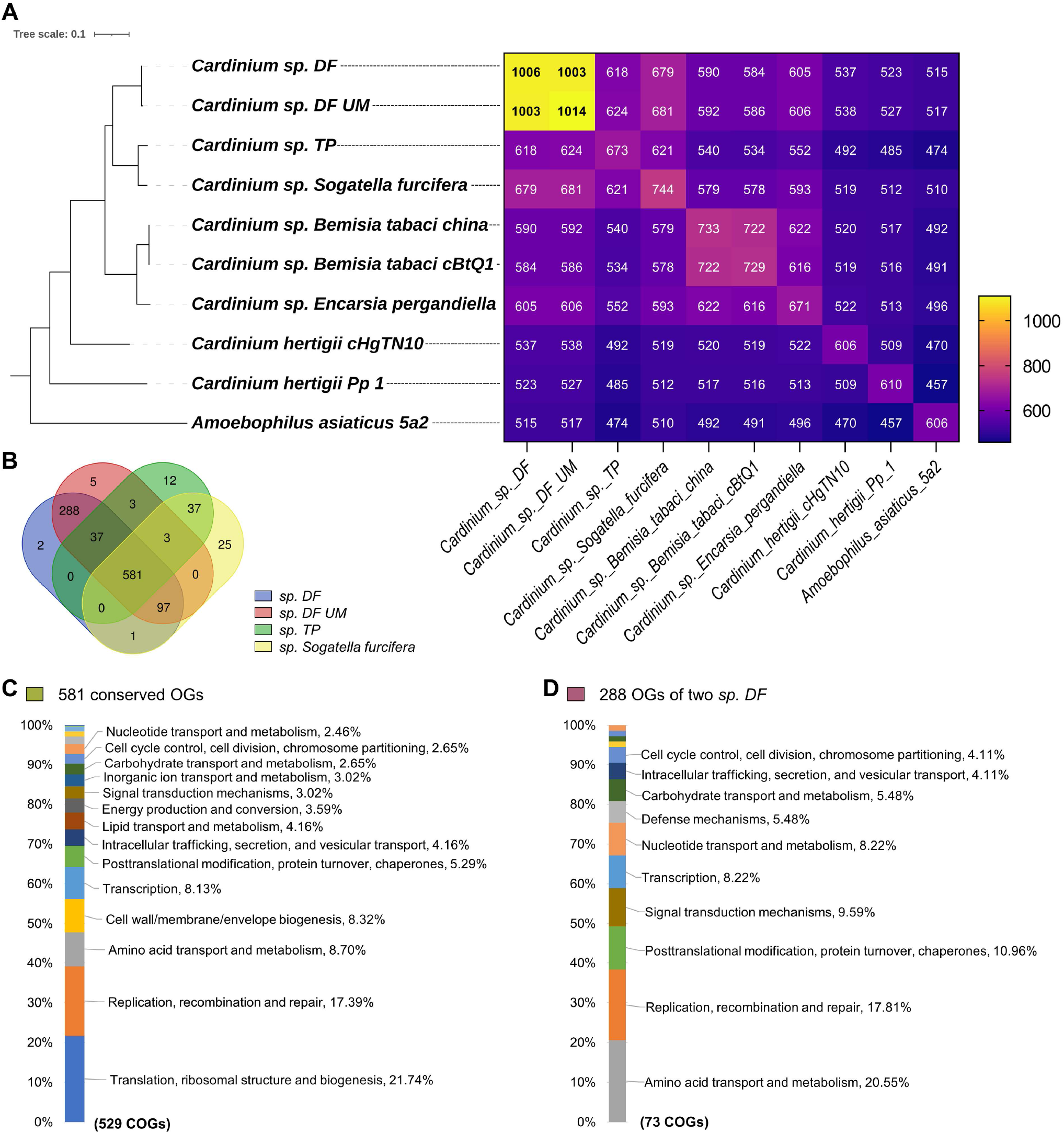
Comparative genomics analysis of *Cardinium* species. **(A)** Matrix of species-to-species overlapped orthogroup number. The protein sequences annotated in the genomes of nine *Cardinium* species and *Amoebophilus asiaticus* as outgroup were comparatively analyzed by the program OrthoFinder and assigned into orthogroups (OGs) according to the sequence similarity The phylogenetic tree was adapted from Fig. S2A. (B) Venn diagram of four *Cardinium* bacteria. The orthogroup lists of the four *Cardinium* genomes were used to generate the Venn diagram, following the analysis of OrthoFinder. **(C)** Functional COGs of the genes of *Cardinium sp. DF* in the 581 conserved OGs among the four *Cardinium* genomes. (D) Functional Clusters of Orthologous Groups (COGs) of the genes of *Cardinium sp. DF* in the 288 conserved OGs specific in the two *Cardinium sp. DF* genomes. The COG category was annotated by eggnog-mapper v2.1.5 and only the functional COGs over >2% were labeled.

Comparative genomics provided an outline for the reductive genome evolution of *Cardinium* species. Two *Cardinium sp. DF* genomes were assigned more OGs than others (Fig. 2A), which is consistent with their relatively larger genome sizes (Table S2). In the Venn diagram of four closely related *Cardinium* species (Fig. 2B), 581 OGs were conserved among the four genomes, and the genes of *Cardinium sp. DF* within these OGs were assigned to 529 functional Clusters of Orthologous Groups (COGs), in which the top two COG categories were “Translation, ribosomal structure and biogenesis” (21.74%) and “Replication, recombination and repair” (17.39%) (Fig. 2C). For the 288 OGs of two *Cardinium sp. DF* not found in the genomes of *Cardinium sp. TP* and *Cardinium sp. Sogatella furcifera*, the genes of *Cardinium sp. DF* were assigned to 73 functional COGs (Fig. 2D), when both the assembly sizes of *Cardinium sp. TP* and *Cardinium sp. Sogatella furcifera* were over 150 Kb smaller than that of the two *Cardinium sp. DF*.

To further explore genome reduction, we focused on the high-quality genome of *Cardinium sp. DF* and combined its two short contigs (Fig. 1B) for further analysis. The sequencing coverage of contigs A and B was estimated to be 146.4 X and 231.0 X, respectively, while that of the chromosomal genome was only 108.6 X (TGS reads in Table S1). The higher coverages of the two short contigs suggest that they are replicable elements in *Cardinium sp. DF*.

In the dot plot of the short contig A of *Cardinium sp. DF* (Fig. S5A), the repeated terminal sequences suggested that it was a circular plasmid containing a pair of inverted repeats (approximately 920 bp). The truncated part (1-31,550 bp) of contig A was considered circular plasmid DNA and named Plasmid A (Fig. S5B). It was annotated with 28 protein-coding genes by Prokka ^14^ (Fig. S5B), in which only three genes were assigned functional names, including two transposase genes and the cell division protein PomZ. More gene annotations were performed by eggNOG-mapper ^15,16^ (Table S3). Except for those recombination-related genes, including relaxase, transposase and resolvase, three NUDIX hydrolase domain-containing genes (GPDKAJLJ_00012-14) were annotated as tandemly arrayed. A SymE toxin gene (GPDKAJLJ_00010) in the type I toxin-antitoxin system was also annotated, and this gene is located in the chromosomal genome of *Escherichia coli* ^17^. To further explore Plasmid A, all encoded proteins were compared against those annotated in its chromosomal genome, and 21 of 28 proteins could be well matched with an E-value cut-off of 1E-6 (Table S4), especially for the Tn3 family transposase TnEc1, GPDKAJLJ_00027 that has multiple highly similar copies in the genome (Table S5).

For the other contig B, the dot plot showed complicated features with a wide range of repeated sequences (Fig. S6A). In the annotation, 8 protein-coding genes were identified as encoding transposases by Prokka ^14^ (Fig. S6B), and more functional annotations were performed by eggNOG-mapper ^15,16^ (Table S6). Contig B was also suggested to be a circular plasmid and called Plasmid B when the two ends could be connected by TGS long reads (Fig. S6C). Concurrent with that in Plasmid A, partial proteins encoded by Plasmid B present high similarities to those in the genome, such as DIOAJDMK_00073 of Plasmid B, which shares 90.9% identity and 87.8% coverage with GPMKIAHG_00283 of the genome (Table S7). Therefore, the two plasmids were suggested to be endogenous plasmids that share homologous genes with the chromosomal genome.

With the two extrachromosomal plasmids (Fig. 1B), a range of unexpected gene conservation (Table S8) and gene synteny alignments (Fig. 3A and B) illuminated the molecular mechanism of the reductive genome evolution of *Cardinium sp. DF*. Genes such as the N-terminal domain of reverse transcriptase DIOAJDMK_00062 of Plasmid B decayed in the genome but remained in Plasmid B of *Cardinium sp. DF* (Table S8).

**Fig. 3.**
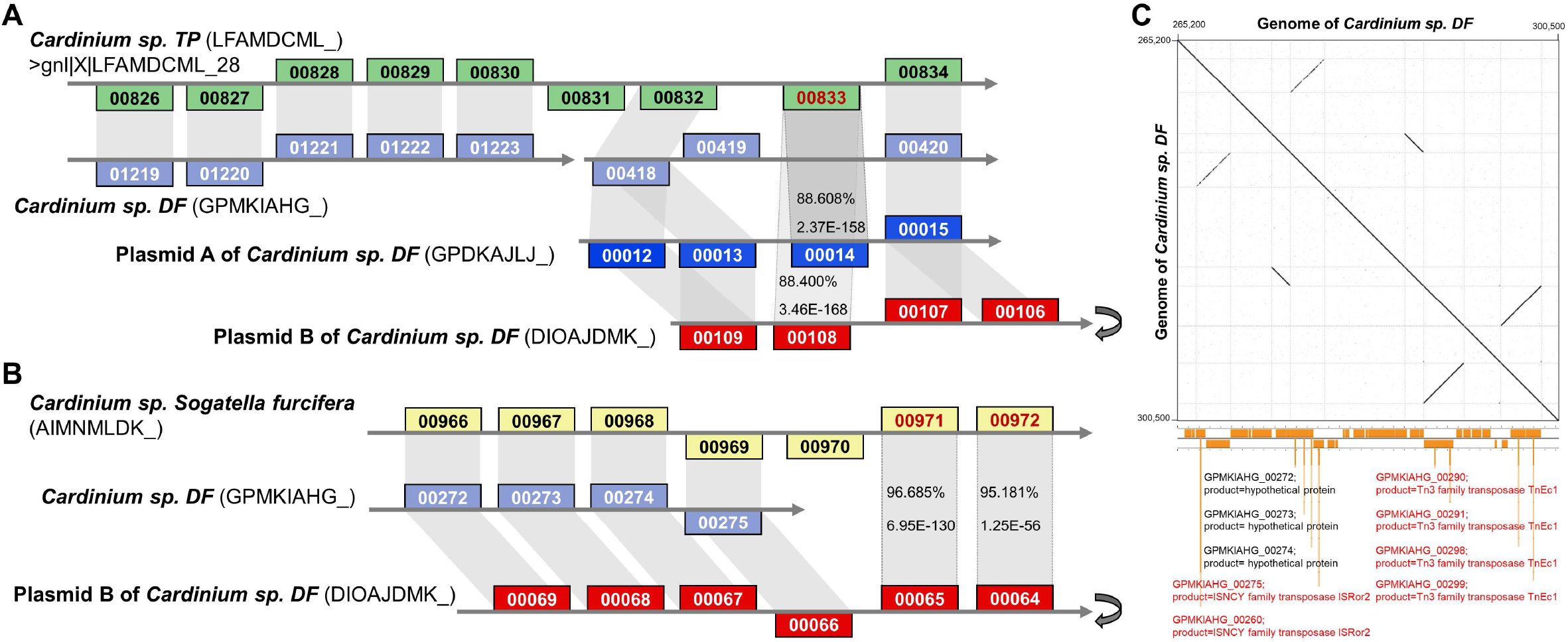
Alignment analysis of the genome reduction in *Cardinium sp. DF*. **(A)** The NUDIX hydrolase gene LFAMDCML_00833 encoded by the contig gnl|X|LFAMDCML_28 of *Cardinium sp. TP* presented high percentages of identities, 88.608% and 88.400% with GPDKAJLJ_0014 of Plasmid A and DIOAJDMK_00108 of Plasmid B respectively; but has no homolog in the genome of *Cardinium sp. DF*. **(B)** The NUBPL iron-transfer P-loop NTPase gene AIMNMLDK_00971 and the hypothetical protein gene AIMNMLDK_00972 of *Cardinium sp. Sogatella furcifera* cannot find homolog in the genome of *Cardinium sp. DF* despite of the conserved synteny in their upstream genes; but shared high similarities with the two genes, DIOAJDMK_00065 and DIOAJDMK_00064 of Plasmid B, respectively. The genes above the gray backbone are located on the plus strand, while those below the backbone are on the minus strand. All homologous genes were connected by the grey bands. The grey turnover symbol means reverse complement. (C) Dot plot of *Cardinium sp. DF* genome region (265200-300500 bp) containing three pairs of repeats. The partial gene annotations were adapted from Fig. 1 A, and three inverted pairs of transposase genes were highlighted in red color. The dot plot was generated by Gepard v2.1.

We selected two reduced loci of *Cardinium sp. DF* to further explore. First, the NUDIX hydrolase gene LFAMDCML_00833 encoded by the contig gnl|X|LFAMDCML_28 of *Cardinium sp. TP* presented an unexpectedly high percentages of identity, 88.608% and 88.400% with GPDKAJLJ_00014 of Plasmid A and DIOAJDMK_00108 of Plasmid B, respectively, but has no homologue in the genome of *Cardinium sp. DF* (Fig. 3A and S7A). Because other genes on the contig gnl|X|LFAMDCML_28 shared high conservation with two regions of the genome of *Cardinium sp. DF* (Fig. 3A), this short contig (12,607 bp) was considered part of the chromosomal genome of *Cardinium sp. TP*. Therefore, we proposed that this NUDIX hydrolase gene has decayed in the genome of *Cardinium sp. DF*, but interestingly, two copies are retained in its plasmids.

Second, the NUBPL iron-transfer P-loop NTPase gene AIMNMLDK_00971 and the hypothetical protein gene AIMNMLDK_00972 of *Cardinium sp. Sogatella furcifera* do not have homologues in the genome of *Cardinium sp. DF*, albeit with the conserved synteny in their upstream genes GPMKIAHG_00272-275, but unexpectedly shared high similarities with the two genes, DIOAJDMK_00065 and DIOAJDMK_00064 of Plasmid B (Fig. 3B, S7B and C). In the homologous location of the genome of *Cardinium sp. DF*, we identified three inverted pairs of highly conserved transposase genes (Fig. 3C), which are possibly related to the molecular mechanism of this reduction. GPMKIAHG_00272-275 are located on a pair of inverted repeats and GPMKIAHG_00275 was annotated as the ISNCY family transposase ISRor2 (Fig. 3C).

Similarly, we identified inverted pairs of resolvase genes in the two plasmids (Fig. S8). The inverted pair of resolvase genes in Plasmid A constitutes a possible transposon similar to those in the Tn-3 family ^18^, along with the downstream transposase gene GPDKAJLJ_00027 (Fig. S8A). Notably, the five genes of Plasmid A, GPDKAJLJ_00023-27 have high-quality homologous genes in the genome of *Cardinium sp. DF* (Fig. S8A, Table S4). In addition, the two resolvase genes DIOAJDMK_00026 and DIOAJDMK_00027 of Plasmid B were decayed in the genome (Table S8). Resolvases often function in a dimer state and can maintain the plasmids in a monomeric state ^19,20^.

One of the most prominent and widely accepted hypotheses, known as Muller’s ratchet, explains the genome reduction as accumulative deletions in the context of asexual reproduction without recombination ^1,2^. In this study, the integrative analyses of the two endogenous plasmids in *Cardinium sp. DF* provide interesting insights into the reductive genome evolution in host-associated bacteria (Fig. S9). To the best of our knowledge, this study introduced endogenous plasmids to the process of genome reduction for the first time. The endogenous plasmids play at least two roles in the genome evolution of *Cardinium sp. DF*. First, they provide homologous genes for the genome and the possibility for recombination (Fig. S9). Second, when genes decay in the chromosomal genome, their homologous genes remaining in the plasmid could still perform necessary functions. In addition, inverted repeats of transposases and resolvases may underpin the molecular process of genome reduction with the aid of endogenous plasmids. Collectively, these endogenous plasmids provide informative snapshots and valuable resources for exploring the ongoing reductive genome evolution of host-associated bacteria.

