## Supplementary Materials for "Endogenous Plasmids and Reductive Genome Evolution in Host-Associated Bacteria"

for

**Supplementary Materials:**

Materials and Methods

Tables S1 to S8

Figures S1 to S9

References

#### Materials and Methods

*Genome assembly and annotation*

All sequencing data used in this study can be found with NCBI accessions in Table S1. To assemble the genome of *Cardinium sp. DF*, the initial genome assembly was constructed with PacBio long reads using Flye v2.6 [1]. Then, further scaffolding was performed by SSPACE Basic v2.0 [2] with paired-end Illumina short reads [3] and SSPACE LongRead v1.1 [4] with PacBio long reads [5] to achieve better continuity. Sequence polishing was finished by Pilon v1.22 [6] with all Illumina short reads. Further scaffolding was performed with SSPACE-LongRead v.1.1 [4] using Oxford Nanopore Technologies (ONT) sequencing reads [7] and gaps were filled with the raw ONT reads using LR_Gapcloser v1.0 [8]. The chromosomal genome and two plasmids located in contigs of of *Cardinium sp. DF* was *de novo* assembled along with the genome of *Dermatophagoides farinae*.

As for the genome of *Cardinium sp. TP*, the assembly was mainly based on NGS short reads (Table S1). NGS reads from three studies were collected for assembling draft genomes using SPAdes v3.13.1 [9] in meta mode respectively. From their assembled sequences, three 16S rRNA sequences of *Cardinium* were identified and confirmed as 100% identities. Then combined NGS reads from the three studies (Table S1) were reassembled using SPAdes v3.13.1 [9] in meta mode. The draft genome was further scaffolded using SSPACE Basic v2.0 [2] with paired-end NGS short reads [3] and SSPACE LongRead v1.1 [4] with PacBio long reads [5] (Table S1). Finally, within the assembled sequences, 33 contigs were assigned as from *Cardinium* genus and considered as the genome assembly of *Cardinium sp. TP*.

The genome annotations were performed by Prokka v1.14.6 [10] and more functional annotations were added by eggNOG-mapper [11,12]. The genome and annotation were visualized by the online tool Proksee (https://proksee.ca/) [13]. To estimate the sequencing coverage, NGS and TGS reads were mapped to the genome by Bowtie2 v2.3.5.1 [14] and Blasr [15], then transformed, sorted and coverage calculated by Samtools v1.9 [16]. Visualization of reads mapping was performed by Integrative Genomics Viewer (IGV) [17].

*Whole genome alignment*

To understand the sequence similarities, whole genome alignment was performed and visualized using AliTV (https://alitvteam.github.io/AliTV/d3/AliTV.html) [18]. Additionally, dot plots were generated by Gepard v2.1 [19] to identify and visualize conserved and repeated regions.

*Comparative genomics*

To explore the evolutionary relationships, comparative genomics analysis was performed among *Cardinium* assemblies (Table S2). All the *Cardinium* genome assemblies were annotated by Prokka v1.14.6 [10] and all the annotated proteomes were assigned into orthogroups (or gene families) based on protein sequence similarities by OrthoFinder v2.5.4 [20]. Then, Venn diagram was performed to identify specific orthogroups using an online tool (https://bioinformatics.psb.ugent.be/webtools/Venn/).

Then phylogenetic analysis was performed based on the sequence alignment of 295 single-copy orthogroups. Firstly, protein sequences in the 295 single-copy orthogroups were extracted and aligned by MAFFT [21], then edited in Gblocks [22] with the option '-t=p' to generate sequence alignment of conserved amino-acid residues. Finally, the sequence alignment was used to construct the phylogenetic tree in maximum likelihood algorithm and 100 bootstrap replicates by RAxML v8.2.12 [23] with the options '-m PROTCATWAG -f a -# 100'. The other phylogenetic tree based on 16S rRNA (Table S2) was constructed by MEGA v11.0.11 [24] with maximum likelihood (ML) algorithm in the JTT (Jones-Taylor-Thornton) model and 100 bootstrap replicates. The phylogenetic trees were finally edited by the online tool Interactive Tree of Life (iTOL, https://itol.embl.de/itol.cgi) [25].

*Genome reduction analysis*

To explore the protein homology, sequence similarity was analyzed by BLASTP v2.9.0 [26] and sequence alignment was performed by the online tool Clustal Omega [27]. Along with the orthogroup assignment, a range of genome reduction loci were identified (Table S8) and further analyzed in gene synteny alignments.

*Data availability*

All the NCBI accessions of sequencing data are in Table S1. The genome assembly and annotation of *Cardinium sp. DF* and *Cardinium sp. TP* have been deposited in the NCBI database under BioProject accessions PRJNA174061 and PRJNA706095 respectively.

### Supplementary tables

#### Table. S1. Genomic sequencing data of two astigmatic mites

NCBI BioProject and SRA accessions of the genomic sequencing data used in this study. NGS, next generation sequencing; TGS, third generation sequencing.

| Species name | NCBI BioProject | NCBI SRA | Sequencing platform | Sequencing type |
|---|---|---|---|---|
| *D. farinae* | PRJNA174061 | SRR3098386 | HiSeq 2000 | NGS |
| | | SRR3098964 | HiSeq 2000 | NGS |
| | | SRR13787846 | PacBio Sequel | TGS |
| | | SRR13741682 | Nanopore GridION | TGS |
| *T. putrescentiae* | PRJNA706095 | SRR13837415 | HiSeq 4000 | NGS |
| | | SRR13837416 | PacBio Sequel | TGS |
| | PRJNA777351 | SRR16709505 | NovaSeq 6000 | NGS |
| | PRJNA656450 | SRR12427973 | NovaSeq 6000 | NGS |

### Table. S2. NCBI GenBank accessions of the sequences used in this study

| Name used in this study | Host species | GenBank accession | Genome size (bp) | Number of contig/scaffold |
|---|---|---|---|---|
| *Cardinium sp. DF* | *Dermatophagoides farinae* | N. A. [a] | 1,259,597 | 1 |
| *Cardinium sp. DF UM* | *Dermatophagoides farinae* | GCA_007559345.1 | 1,258,868 | 1 |
| *Cardinium sp. TP* | *Tyrophagus putrescentiae* | N. A. [a] | 914,750 | 33 |
| Cardinium sp. *Sogatella furcifera* | *Sogatella furcifera* | GCA_003351905.1 | 1,103,593 | 1 |
| *Cardinium sp. Bemisia tabaci china* | *Bemisia tabaci* | GCA_004300865.1 | 1,012,588 | 3 |
| *Cardinium sp. Bemisia tabaci cBtQ1* | *Bemisia tabaci* | GCA_000689375.1 | 996,809 | 11 |
| *Cardinium sp. Encarsia pergandiella* | *Encarsia pergandiella* | GCA_000304455.1 | 944,930 | 2 |
| *Cardinium hertigii cHgTN10* | *Heterodera glycines* | GCA_003176915.1 | 1,193,042 | 1 |
| *Cardinium hertigii Pp_1* | *Pratylenchus penetrans* | GCA_003788695.1 | 1,358,212 | 27 |
| *Amoebophilus asiaticus 5a2* | *Acanthamoeba sp. TUMSJ-321* | GCA_000020565.1 | 1,884,364 | 1 |
| *Cardinium sp. Oppiella nova* | *Oppiella nova* | AY279414 [b] | | |
| *Cardinium sp. Achiperia coleoptrata* | *Achiperia coleoptrata* | MG889457 [b] | | |
| *Cardinium sp. Microzetorchestes emeryi -1* | *Microzetorchestes emeryi* | MG889458 [b] | | |
| *Cardinium sp. Microzetorchestes emeryi -2* | *Microzetorchestes emeryi* | MG889459 [b] | | |

a Not applicable. No NCBI GenBank accession assigned yet.
b Partial 16s rRNA sequences.

#### Table. S3. EggNOG-mapper output of protein sequences of Plasmid A

The annotated protein sequences of Plasmid A (Fig. S5) were annotated by eggnog-mapper v2.1.5.

| #query | Description | PFAMs | Preferred_name |
|---|---|---|---|
| GPDKAJLJ_00002 | Psort location Cytoplasmic, score 8.96 | AAA-ATPase_like,PDDEXK_9 | - |
| GPDKAJLJ_00003 | TraM recognition site of TraD and TraG | T4SS-DNA_transf,TraG-D_C,TrwB_AAD_bind | - |
| GPDKAJLJ_00004 | Relaxase/Mobilisation nuclease domain | Relaxase | bmgA |
| GPDKAJLJ_00006 | - | - | - |
| GPDKAJLJ_00007 | putative transposase, YhgA-like | Transposase_31 | - |
| GPDKAJLJ_00008 | - | - | - |
| GPDKAJLJ_00009 | AAA domain | AAA_31 | - |
| GPDKAJLJ_00010 | toxin SymE, type I toxin-antitoxin system | SymE_toxin | - |
| GPDKAJLJ_00011 | helix-turn-helix XRE-family like proteins | HTH_3 | - |
| GPDKAJLJ_00012 | GDP-mannose mannosyl hydrolase activity | NUDIX | - |
| GPDKAJLJ_00013 | GDP-mannose mannosyl hydrolase activity | NUDIX | - |
| GPDKAJLJ_00014 | NUDIX hydrolase | NUDIX | - |
| GPDKAJLJ_00015 | symporter activity | HD_4,HTH_18,SSF | - |
| GPDKAJLJ_00023 | Resolvase | Resolvase | - |
| GPDKAJLJ_00024 | - | AAA-ATPase_like,HTH_3,PDDEXK_9 | - |
| GPDKAJLJ_00025 | P-loop ATPase and inactivated | VirE | virE |
| GPDKAJLJ_00026 | Resolvase | Resolvase | - |
| GPDKAJLJ_00027 | Tn3 transposase DDE domain | DDE_Tnp_Tn3,DUF4158 | - |
| GPDKAJLJ_00028 | P-loop ATPase and inactivated | VirE | virE |

#### Table. S4. BLASTP output of annotated protein sequences of Plasmid A

The annotated protein sequences of Plasmid A (Fig. S5) were mapped to those of the genome of *Cardinium sp. DF* (Fig. 1A) using BLASTP with options "-evalue 1e-6 -outfmt 6 -max_target_seqs 1" and then sorted by bit score.

| query id | subject id | % identity | alignment length | mismatches | gap opens | q. start | q. end | s. start | s. end | evalue | bit score |
|---|---|---|---|---|---|---|---|---|---|---|---|
| GPDKAJLJ_00027 | GPMKIAHG_00026 | 100 | 818 | 0 | 0 | 1 | 818 | 88 | 905 | 0 | 1691 |
| GPDKAJLJ_00015 | GPMKIAHG_00420 | 70.766 | 992 | 284 | 2 | 3 | 993 | 4 | 990 | 0 | 1415 |
| GPDKAJLJ_00024 | GPMKIAHG_00626 | 77.304 | 586 | 133 | 0 | 1 | 586 | 1 | 586 | 0 | 942 |
| GPDKAJLJ_00002 | GPMKIAHG_00240 | 76.867 | 549 | 116 | 2 | 1 | 538 | 1 | 549 | 0 | 871 |
| GPDKAJLJ_00028 | GPMKIAHG_00251 | 62.702 | 496 | 181 | 3 | 1 | 494 | 1 | 494 | 0 | 629 |
| GPDKAJLJ_00007 | GPMKIAHG_00518 | 82.555 | 321 | 44 | 2 | 7 | 315 | 3 | 323 | 0 | 542 |
| GPDKAJLJ_00026 | GPMKIAHG_00891 | 100 | 219 | 0 | 0 | 1 | 219 | 1 | 219 | 1.76E-167 | 455 |
| GPDKAJLJ_00023 | GPMKIAHG_00834 | 99.543 | 219 | 1 | 0 | 1 | 219 | 1 | 219 | 3.56E-166 | 452 |
| GPDKAJLJ_00022 | GPMKIAHG_00003 | 57.25 | 400 | 158 | 6 | 22 | 411 | 1 | 397 | 2.39E-146 | 417 |
| GPDKAJLJ_00025 | GPMKIAHG_00293 | 64.583 | 192 | 67 | 1 | 1 | 191 | 38 | 229 | 3.54E-83 | 241 |
| GPDKAJLJ_00003 | GPMKIAHG_01234 | 29.893 | 562 | 341 | 15 | 20 | 551 | 81 | 619 | 1.01E-65 | 227 |
| GPDKAJLJ_00021 | GPMKIAHG_00253 | 45.802 | 262 | 116 | 7 | 4 | 242 | 13 | 271 | 5.49E-69 | 209 |
| GPDKAJLJ_00012 | GPMKIAHG_00418 | 83.673 | 98 | 16 | 0 | 1 | 98 | 47 | 144 | 6.06E-58 | 170 |
| GPDKAJLJ_00019 | GPMKIAHG_00295 | 86.364 | 44 | 6 | 0 | 1 | 44 | 145 | 188 | 4.04E-23 | 82.4 |
| GPDKAJLJ_00013 | GPMKIAHG_00418 | 81.818 | 44 | 8 | 0 | 1 | 44 | 1 | 44 | 4.97E-24 | 82.4 |
| GPDKAJLJ_00020 | GPMKIAHG_00295 | 48.837 | 86 | 39 | 1 | 3 | 88 | 14 | 94 | 5.08E-20 | 76.3 |
| GPDKAJLJ_00009 | GPMKIAHG_00596 | 27.602 | 221 | 146 | 6 | 58 | 272 | 3 | 215 | 1.44E-17 | 76.3 |
| GPDKAJLJ_00022 | GPMKIAHG_00003 | 32.609 | 184 | 103 | 7 | 67 | 233 | 220 | 399 | 6.87E-16 | 73.9 |
| GPDKAJLJ_00001 | GPMKIAHG_01231 | 37.5 | 88 | 55 | 0 | 12 | 99 | 14 | 101 | 3.04E-18 | 69.3 |
| GPDKAJLJ_00018 | GPMKIAHG_00295 | 60.938 | 64 | 25 | 0 | 12 | 75 | 187 | 250 | 7.26E-16 | 64.7 |
| GPDKAJLJ_00016 | GPMKIAHG_00162 | 39.655 | 58 | 34 | 1 | 14 | 71 | 210 | 266 | 9.38E-07 | 39.7 |

#### Table. S5. BLASTP output of GPDKAJLJ_00027

The Tn3 family transposase TnEc1, GPDKAJLJ_00027 of Plasmid B (Fig. S6) was mapped to those of the genome of *Cardinium sp. DF* (Fig. 1A) using BLASTP with options "-evalue 1e-6 -outfmt 6 -max_target_seqs 1".

| query id | subject id | % identity | alignment length | mismatches | gap opens | q. start | q. end | s. start | s. end | evalue | bit score |
|---|---|---|---|---|---|---|---|---|---|---|---|
| GPDKAJLJ_00027 | GPMKIAHG_00026 | 100 | 818 | 0 | 0 | 1 | 818 | 88 | 905 | 0 | 1691 |
| GPDKAJLJ_00027 | GPMKIAHG_00833 | 99.51 | 818 | 4 | 0 | 1 | 818 | 88 | 905 | 0 | 1680 |
| GPDKAJLJ_00027 | GPMKIAHG_00658 | 97.99 | 797 | 16 | 0 | 22 | 818 | 1 | 797 | 0 | 1610 |
| GPDKAJLJ_00027 | GPMKIAHG_00290 | 99.72 | 707 | 2 | 0 | 112 | 818 | 1 | 707 | 0 | 1458 |
| GPDKAJLJ_00027 | GPMKIAHG_01011 | 99.55 | 664 | 3 | 0 | 155 | 818 | 1 | 664 | 0 | 1372 |
| GPDKAJLJ_00027 | GPMKIAHG_00299 | 98.85 | 435 | 5 | 0 | 384 | 818 | 1 | 435 | 0 | 894 |
| GPDKAJLJ_00027 | GPMKIAHG_00627 | 98.55 | 414 | 6 | 0 | 405 | 818 | 1 | 414 | 0 | 851 |
| GPDKAJLJ_00027 | GPMKIAHG_00893 | 99.75 | 394 | 1 | 0 | 1 | 394 | 1 | 394 | 0 | 809 |
| GPDKAJLJ_00027 | GPMKIAHG_00628 | 98.24 | 397 | 7 | 0 | 1 | 397 | 88 | 484 | 0 | 803 |
| GPDKAJLJ_00027 | GPMKIAHG_00298 | 97.87 | 376 | 8 | 0 | 1 | 376 | 88 | 463 | 0 | 759 |
| GPDKAJLJ_00027 | GPMKIAHG_00894 | 98.72 | 235 | 3 | 0 | 584 | 818 | 45 | 279 | 2.00E-165 | 481 |
| GPDKAJLJ_00027 | GPMKIAHG_00894 | 59.78 | 92 | 31 | 2 | 393 | 484 | 2 | 87 | 4.00E-22 | 94 |
| GPDKAJLJ_00027 | GPMKIAHG_01113 | 98.14 | 161 | 3 | 0 | 1 | 161 | 150 | 310 | 7.00E-104 | 322 |
| GPDKAJLJ_00027 | GPMKIAHG_01110 | 98.14 | 161 | 3 | 0 | 1 | 161 | 150 | 310 | 1.00E-103 | 322 |
| GPDKAJLJ_00027 | GPMKIAHG_01012 | 94.74 | 133 | 7 | 0 | 20 | 152 | 3 | 135 | 5.00E-79 | 249 |
| GPDKAJLJ_00027 | GPMKIAHG_00076 | 94.57 | 129 | 5 | 1 | 1 | 127 | 88 | 216 | 2.00E-76 | 246 |
| GPDKAJLJ_00027 | GPMKIAHG_00291 | 100 | 109 | 0 | 0 | 1 | 109 | 88 | 196 | 5.00E-69 | 224 |
| GPDKAJLJ_00027 | GPMKIAHG_00010 | 36.15 | 296 | 174 | 7 | 424 | 710 | 1 | 290 | 2.00E-49 | 173 |
| GPDKAJLJ_00027 | GPMKIAHG_00011 | 25.41 | 362 | 258 | 6 | 9 | 360 | 15 | 374 | 3.00E-29 | 116 |
| GPDKAJLJ_00027 | GPMKIAHG_00009 | 37.14 | 105 | 66 | 0 | 711 | 815 | 12 | 116 | 1.00E-17 | 76.6 |
| GPDKAJLJ_00027 | GPMKIAHG_01013 | 92.59 | 27 | 2 | 0 | 1 | 27 | 88 | 114 | 2.00E-10 | 55.8 |

### Table. S6. EggNOG-mapper output of protein sequences of Plasmid B

The annotated protein sequences of Plasmid B (Fig. S6) were annotated by eggnog-mapper v2.1.5.

| #query | Description | PFAMs | Preferred_name |
|---|---|---|---|
| DIOAJDMK_00001 | DNA primase activity | DUF3991,DnaB_C,Toprim_2,Toprim_3,zf-CHC2 | - |
| DIOAJDMK_00002 | DNA primase activity | DUF3991,DnaB_C,Toprim_2,Toprim_3,zf-CHC2 | - |
| DIOAJDMK_00004 | DNA primase activity | PriCT_2,Prim-Pol,Toprim_2,Toprim_3,zf-CHC2 | - |
| DIOAJDMK_00006 | plasmid recombination enzyme | Mob_Pre | - |
| DIOAJDMK_00009 | TraM recognition site of TraD and TraG | T4SS-DNA_transf,TraG-D_C,TrwB_AAD_bind | - |
| DIOAJDMK_00010 | response to abiotic stimulus | Ank_2,Ank_3,Ank_4,Ank_5,Rhodanese | - |
| DIOAJDMK_00016 | type IV secretory pathway VirB4 | DUF3875,DUF87 | - |
| DIOAJDMK_00017 | - | - | - |
| DIOAJDMK_00018 | conjugation | VirB8 | trbF |
| DIOAJDMK_00021 | conjugal transfer protein TraM | Transposon_TraM | - |
| DIOAJDMK_00025 | putative transposase, YhgA-like | Transposase_31 | - |
| DIOAJDMK_00026 | Resolvase, N terminal domain | HTH_7,Resolvase | - |
| DIOAJDMK_00027 | Resolvase, N terminal domain | HTH_7,Resolvase | - |
| DIOAJDMK_00028 | N-acetylmuramoyl-L-alanine amidase | Amidase_2,PG_binding_1 | amiD |
| DIOAJDMK_00029 | NLPC_P60 stabilising domain, N term | NLPC_P60,N_NLPC_P60,SH3_6,SH3_7 | - |
| DIOAJDMK_00030 | symporter activity | HD_4,HTH_18,SSF | - |
| DIOAJDMK_00031 | symporter activity | HD_4,HTH_18,SSF | - |
| DIOAJDMK_00032 | putative transposase, YhgA-like | Transposase_31 | - |
| DIOAJDMK_00035 | TIGRFAM Bacteroides conjugative transposon TraM protein | Transposon_TraM | - |
| DIOAJDMK_00037 | conjugation | VirB8 | trbF |
| DIOAJDMK_00038 | - | - | - |
| DIOAJDMK_00040 | type IV secretory pathway VirB4 | DUF3875,DUF87 | - |
| DIOAJDMK_00041 | type IV secretory pathway VirB4 | DUF3875,DUF87 | - |
| DIOAJDMK_00048 | response to abiotic stimulus | Ank_2,Ank_3,Ank_4,Ank_5,Rhodanese | - |
| DIOAJDMK_00049 | response to abiotic stimulus | Ank_2,Ank_3,Ank_4,Ank_5,Rhodanese | - |
| DIOAJDMK_00051 | TraM recognition site of TraD and TraG | T4SS-DNA_transf,TraG-D_C,TrwB_AAD_bind | - |
| DIOAJDMK_00052 | TraM recognition site of TraD and TraG | T4SS-DNA_transf,TraG-D_C,TrwB_AAD_bind | - |
| DIOAJDMK_00054 | plasmid recombination enzyme | Mob_Pre | - |
| DIOAJDMK_00056 | P-loop ATPase and inactivated | VirE | virE |
| DIOAJDMK_00057 | P-loop ATPase and inactivated | VirE | virE |
| DIOAJDMK_00060 | Transposase IS66 family | DDE_Tnp_IS66,zf-IS66 | - |
| DIOAJDMK_00061 | to TIGR01784 | PDDEXK_2 | - |
| DIOAJDMK_00062 | N-terminal domain of reverse transcriptase | GIIM,RVT_1,RVT_N | - |
| DIOAJDMK_00063 | belongs to the 'phage' integrase family | Phage_int_SAM_5,Phage_integrase | - |
| DIOAJDMK_00065 | NUBPL iron-transfer P-loop NTPase | AAA_31,CbiA | - |
| DIOAJDMK_00066 | putative transposase, YhgA-like | Transposase_31 | - |
| DIOAJDMK_00069 | conjugal transfer protein TraM | Transposon_TraM | - |
| DIOAJDMK_00071 | conjugation | VirB8 | trbF |
| DIOAJDMK_00072 | - | - | - |
| DIOAJDMK_00073 | type IV secretory pathway VirB4 | DUF3875,DUF87 | - |
| DIOAJDMK_00078 | response to abiotic stimulus | Ank_2,Ank_3,Ank_4,Ank_5,Rhodanese | - |
| DIOAJDMK_00079 | TraM recognition site of TraD and TraG | T4SS-DNA_transf,TraG-D_C,TrwB_AAD_bind | - |
| DIOAJDMK_00082 | plasmid recombination enzyme | Mob_Pre | - |
| DIOAJDMK_00087 | DNA primase activity | PriCT_2,Prim-Pol,Toprim_2,Toprim_3,zf-CHC2 | - |
| DIOAJDMK_00088 | DNA primase activity | DUF3991,DnaB_C,Toprim_2,Toprim_3,zf-CHC2 | - |
| DIOAJDMK_00089 | P-loop ATPase and inactivated | VirE | virE |
| DIOAJDMK_00090 | P-loop ATPase and inactivated | VirE | virE |
| DIOAJDMK_00091 | cellulose biosynthesis protein BcsQ | AAA_31 | - |
| DIOAJDMK_00092 | ParB-like nuclease domain | ParBc | - |
| DIOAJDMK_00093 | GDP-mannose mannosyl hydrolase activity | NUDIX | - |
| DIOAJDMK_00094 | domain of unknown function (DUF4277) | DDE_Tnp_1,DUF4277 | - |
| DIOAJDMK_00095 | winged helix-turn helix | HTH_29,rve,rve_3 | - |
| DIOAJDMK_00096 | winged helix-turn helix | HTH_29,rve,rve_3 | - |
| DIOAJDMK_00097 | Transposase IS66 family | DDE_Tnp_IS66,zf-IS66 | - |
| DIOAJDMK_00098 | - | - | - |
| DIOAJDMK_00103 | response to abiotic stimulus | Ank,Ank_2,Ank_3,Ank_4,Ank_5 | - |
| DIOAJDMK_00106 | symporter activity | HD_4,HTH_18,SSF | - |
| DIOAJDMK_00107 | symporter activity | HD_4,HTH_18,SSF | - |
| DIOAJDMK_00108 | NUDIX hydrolase | NUDIX | - |
| DIOAJDMK_00109 | GDP-mannose mannosyl hydrolase activity | NUDIX | - |
| DIOAJDMK_00111 | ATPase MipZ | AAA_31,MipZ | - |
| DIOAJDMK_00113 | P-loop ATPase and inactivated | VirE | virE |
| DIOAJDMK_00114 | DNA primase activity | PriCT_2,Prim-Pol,Toprim_2,Toprim_3,zf-CHC2 | - |
| DIOAJDMK_00116 | plasmid recombination enzyme | Mob_Pre | - |
| DIOAJDMK_00118 | TraM recognition site of TraD and TraG | T4SS-DNA_transf,TraG-D_C,TrwB_AAD_bind | - |
| DIOAJDMK_00119 | response to abiotic stimulus | Ank_2,Ank_3,Ank_4,Ank_5,Rhodanese | - |
| DIOAJDMK_00124 | Resolvase | Resolvase | - |
| DIOAJDMK_00125 | domain of unknown function (DUF4158) | DDE_Tnp_Tn3,DUF4158 | - |
| DIOAJDMK_00126 | Tn3 transposase DDE domain | DDE_Tnp_Tn3,DUF4158 | - |
| DIOAJDMK_00128 | PD-(D/E)XK nuclease family transposase | PDDEXK_2 | - |
| DIOAJDMK_00129 | plasmid maintenance | AAA_31,MipZ | - |
| DIOAJDMK_00130 | Resolvase | Resolvase | - |

### Table. S7. BLASTP output of annotated protein sequences of Plasmid B

The annotated protein sequences of Plasmid B (Fig. S6) were mapped to those of the genome of *Cardinium sp. DF* (Fig. 1A) using BLASTP with options "-evalue 1e-6 -outfmt 6 -max_target_seqs 1" (only those below 1E-100 are shown) and sorted by bit score.

| query id | subject id | % identity | alignment length | mismatches | gap opens | q. start | q. end | s. start | s. end | evalue | bit score |
|---|---|---|---|---|---|---|---|---|---|---|---|
| DIOAJDMK_00073 | GPMKIAHG_00283 | 90.858 | 711 | 65 | 0 | 1 | 711 | 1 | 711 | 0 | 1343 |
| DIOAJDMK_00016 | GPMKIAHG_00283 | 90.577 | 711 | 67 | 0 | 1 | 711 | 1 | 711 | 0 | 1335 |
| DIOAJDMK_00116 | GPMKIAHG_01232 | 89.342 | 638 | 68 | 0 | 1 | 638 | 1 | 638 | 0 | 1184 |
| DIOAJDMK_00054 | GPMKIAHG_01232 | 89.342 | 638 | 68 | 0 | 1 | 638 | 1 | 638 | 0 | 1184 |
| DIOAJDMK_00030 | GPMKIAHG_00375 | 57.352 | 1027 | 429 | 6 | 10 | 1032 | 10 | 1031 | 0 | 1161 |
| DIOAJDMK_00118 | GPMKIAHG_01234 | 92.845 | 601 | 43 | 0 | 17 | 617 | 17 | 617 | 0 | 1154 |
| DIOAJDMK_00079 | GPMKIAHG_01234 | 92.845 | 601 | 43 | 0 | 17 | 617 | 17 | 617 | 0 | 1154 |
| DIOAJDMK_00006 | GPMKIAHG_01232 | 86.614 | 635 | 85 | 0 | 1 | 635 | 1 | 635 | 0 | 1151 |
| DIOAJDMK_00009 | GPMKIAHG_01234 | 92.487 | 599 | 45 | 0 | 1 | 599 | 19 | 617 | 0 | 1146 |
| DIOAJDMK_00010 | GPMKIAHG_00267 | 89.949 | 587 | 59 | 0 | 1 | 587 | 1 | 587 | 0 | 1090 |
| DIOAJDMK_00119 | GPMKIAHG_01234 | 84.588 | 558 | 80 | 2 | 60 | 614 | 597 | 1151 | 0 | 966 |
| DIOAJDMK_00078 | GPMKIAHG_01234 | 84.588 | 558 | 80 | 2 | 60 | 614 | 597 | 1151 | 0 | 966 |
| DIOAJDMK_00056 | GPMKIAHG_00251 | 84.404 | 545 | 85 | 0 | 47 | 591 | 1 | 545 | 0 | 956 |
| DIOAJDMK_00106 | GPMKIAHG_00420 | 75.702 | 605 | 146 | 1 | 3 | 606 | 386 | 990 | 0 | 933 |
| DIOAJDMK_00113 | GPMKIAHG_00021 | 85.857 | 502 | 71 | 0 | 84 | 585 | 1 | 502 | 0 | 870 |
| DIOAJDMK_00041 | GPMKIAHG_00283 | 88.032 | 376 | 45 | 0 | 1 | 376 | 1 | 376 | 0 | 696 |
| DIOAJDMK_00049 | GPMKIAHG_00267 | 83.815 | 346 | 56 | 0 | 1 | 346 | 130 | 475 | 0 | 608 |
| DIOAJDMK_00034 | GPMKIAHG_00273 | 96.382 | 304 | 11 | 0 | 1 | 304 | 1 | 304 | 0 | 584 |
| DIOAJDMK_00022 | GPMKIAHG_00273 | 94.737 | 304 | 16 | 0 | 1 | 304 | 1 | 304 | 0 | 577 |
| DIOAJDMK_00068 | GPMKIAHG_00273 | 92.763 | 304 | 22 | 0 | 1 | 304 | 1 | 304 | 0 | 567 |
| DIOAJDMK_00051 | GPMKIAHG_01234 | 92.491 | 293 | 22 | 0 | 1 | 293 | 323 | 615 | 0 | 558 |
| DIOAJDMK_00076 | GPMKIAHG_01236 | 95.941 | 271 | 11 | 0 | 5 | 275 | 1 | 271 | 0 | 553 |
| DIOAJDMK_00121 | GPMKIAHG_01236 | 95.941 | 271 | 11 | 0 | 5 | 275 | 1 | 271 | 0 | 549 |
| DIOAJDMK_00032 | GPMKIAHG_00275 | 80.665 | 331 | 52 | 1 | 1 | 331 | 1 | 319 | 0 | 546 |
| DIOAJDMK_00066 | GPMKIAHG_00518 | 80.122 | 327 | 53 | 2 | 5 | 323 | 1 | 323 | 0 | 533 |
| DIOAJDMK_00025 | GPMKIAHG_00401 | 79.817 | 327 | 54 | 2 | 1 | 327 | 1 | 315 | 0 | 531 |
| DIOAJDMK_00021 | GPMKIAHG_00272 | 92.708 | 288 | 21 | 0 | 1 | 288 | 1 | 288 | 0 | 516 |
| DIOAJDMK_00040 | GPMKIAHG_00283 | 91.176 | 272 | 24 | 0 | 1 | 272 | 447 | 718 | 1.8E-180 | 514 |
| DIOAJDMK_00035 | GPMKIAHG_00272 | 88.542 | 288 | 31 | 1 | 1 | 286 | 1 | 288 | 5.3E-179 | 490 |
| DIOAJDMK_00069 | GPMKIAHG_00272 | 87.241 | 290 | 35 | 1 | 1 | 290 | 1 | 288 | 1.4E-178 | 489 |
| DIOAJDMK_00017 | GPMKIAHG_00285 | 93.727 | 271 | 17 | 0 | 1 | 271 | 1 | 271 | 1.3E-169 | 465 |
| DIOAJDMK_00072 | GPMKIAHG_00285 | 92.251 | 271 | 21 | 0 | 1 | 271 | 1 | 271 | 2.4E-165 | 454 |
| DIOAJDMK_00124 | GPMKIAHG_00834 | 98.174 | 219 | 4 | 0 | 1 | 219 | 1 | 219 | 2.3E-165 | 450 |
| DIOAJDMK_00088 | GPMKIAHG_00251 | 70.508 | 295 | 87 | 0 | 1 | 295 | 52 | 346 | 3.2E-151 | 430 |
| DIOAJDMK_00107 | GPMKIAHG_00420 | 61.429 | 350 | 129 | 2 | 1 | 349 | 40 | 384 | 5.6E-141 | 420 |
| DIOAJDMK_00012 | GPMKIAHG_01236 | 86.486 | 222 | 30 | 0 | 5 | 226 | 1 | 222 | 1.7E-145 | 402 |
| DIOAJDMK_00125 | GPMKIAHG_01113 | 97.98 | 198 | 4 | 0 | 1 | 198 | 1 | 198 | 3.1E-143 | 398 |
| DIOAJDMK_00018 | GPMKIAHG_00690 | 81.735 | 219 | 40 | 0 | 1 | 219 | 1 | 219 | 1E-137 | 380 |
| DIOAJDMK_00013 | GPMKIAHG_01237 | 82.028 | 217 | 39 | 0 | 46 | 262 | 7 | 223 | 5.3E-131 | 380 |
| DIOAJDMK_00037 | GPMKIAHG_00690 | 85.096 | 208 | 31 | 0 | 13 | 220 | 12 | 219 | 5.2E-137 | 379 |
| DIOAJDMK_00071 | GPMKIAHG_00690 | 85.096 | 208 | 31 | 0 | 13 | 220 | 12 | 219 | 1.8E-136 | 377 |
| DIOAJDMK_00044 | GPMKIAHG_01237 | 83.105 | 219 | 37 | 0 | 181 | 399 | 7 | 225 | 1.9E-131 | 376 |
| DIOAJDMK_00052 | GPMKIAHG_00265 | 84.762 | 210 | 32 | 0 | 8 | 217 | 93 | 302 | 2.9E-129 | 370 |
| DIOAJDMK_00077 | GPMKIAHG_01235 | 94.22 | 173 | 10 | 0 | 1 | 173 | 1 | 173 | 3.9E-123 | 339 |
| DIOAJDMK_00120 | GPMKIAHG_01235 | 94.22 | 173 | 10 | 0 | 1 | 173 | 1 | 173 | 1.2E-122 | 338 |
| DIOAJDMK_00011 | GPMKIAHG_01235 | 91.329 | 173 | 15 | 0 | 1 | 173 | 1 | 173 | 9.7E-121 | 333 |
| DIOAJDMK_00089 | GPMKIAHG_00021 | 82.162 | 185 | 33 | 0 | 1 | 185 | 238 | 422 | 4.5E-111 | 321 |
| DIOAJDMK_00095 | GPMKIAHG_00002 | 95.57 | 158 | 7 | 0 | 1 | 158 | 1 | 158 | 1.6E-108 | 309 |

#### **Table. S8. Candidate genes in the reductive evolution of *Cardinium sp. DF***

These candidate genes encoded by Plasmid A (Fig. S5), or Plasmid B (Fig. S6) shared high percentage of identities with genes encoded by the genome of *Cardinium sp. TP* or *Cardinium sp. Sogatella furcifera* but have no homolog (E-value cut-off: 1E-6) from encoded genes of the genome of *Cardinium sp. DF*. The gene description was annotated by eggnog-mapper v2.1.5.

| **Gene ID** | **Location** | **Description** | **Best-match gene** | **% identity** | **E-value** |
|---|---|---|---|---|---|
| GPDKAJLJ_00014 | Plasmid A | NUDIX hydrolase | LFAMDCML_00833 *Cardinium sp. TP* | 88.608% | 2.37E-158 |
| DIOAJDMK_00026 | Plasmid B | Resolvase, N terminal domain | AIMNMLDK_00049 *Cardinium sp. Sogatella furcifera* | 90.957% | 5.31E-117 |
| DIOAJDMK_00027 | Plasmid B | Resolvase, N terminal domain | AIMNMLDK_00049 *Cardinium sp. Sogatella furcifera* | 91.667% | 8.10E-121 |
| DIOAJDMK_00062 | Plasmid B | N-terminal domain of reverse transcriptase | AIMNMLDK_00186 *Cardinium sp. Sogatella furcifera* | 38.889% | 7.86E-71 |
| DIOAJDMK_00064 | Plasmid B | Hypothetical protein | AIMNMLDK_00972 *Cardinium sp. Sogatella furcifera* | 95.181% | 1.25E-56 |
| DIOAJDMK_00065 | Plasmid B | NUBPL iron-transfer P-loop NTPase | AIMNMLDK_00971 *Cardinium sp. Sogatella furcifera* | 96.685% | 6.95E-130 |
| DIOAJDMK_00108 | Plasmid B | NUDIX hydrolase | LFAMDCML_00833 *Cardinium sp. TP* | 88.400% | 3.46E-168 |
| DIOAJDMK_00131 | Plasmid B | Hypothetical protein | AIMNMLDK_00978 *Cardinium sp. Sogatella furcifera* | 56.574% | 4.06E-94 |
| DIOAJDMK_00134 | Plasmid B | Hypothetical protein | AIMNMLDK_00979 *Cardinium sp. Sogatella furcifera* | 61.905% | 3.11E-34 |

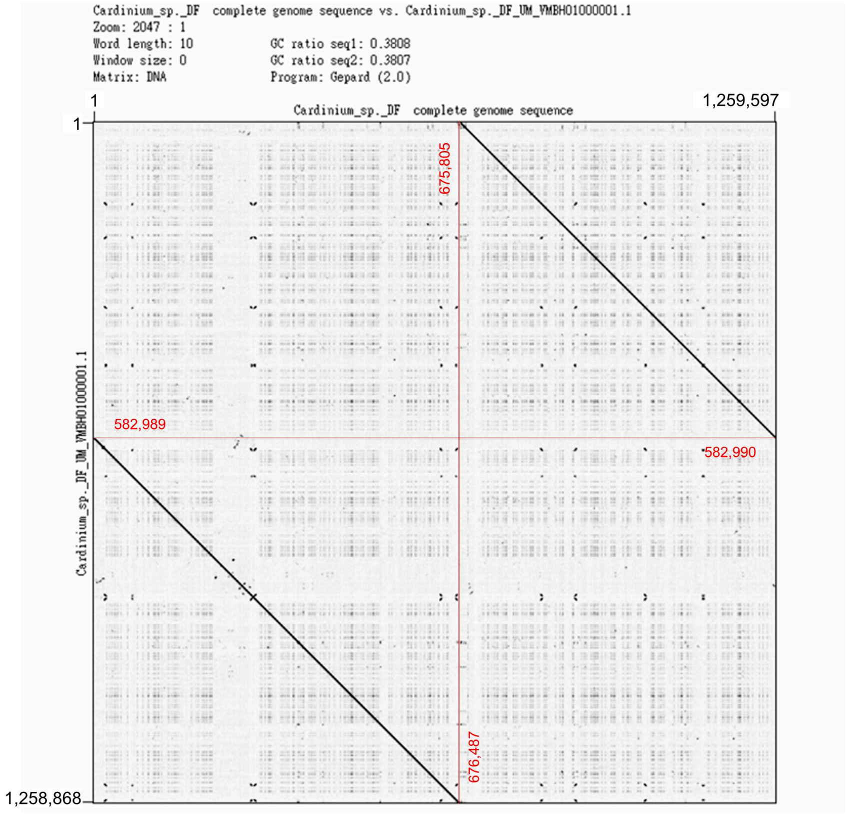

**Fig. S1. Dot plot of two *Cardinium sp. DF* genomes**

The dot plot was generated by Gepard v2.1 with the genome of *Cardinium sp. DF* and the longest contig of *Cardinium sp. DF UM*. The key junction sites were highlighted and labeled. The staggered feature between two genomes suggested both were circular DNA and the region 675,805-676,487 of *Cardinium sp. DF* has not match in *Cardinium sp. DF UM*.

### Supplementary figures

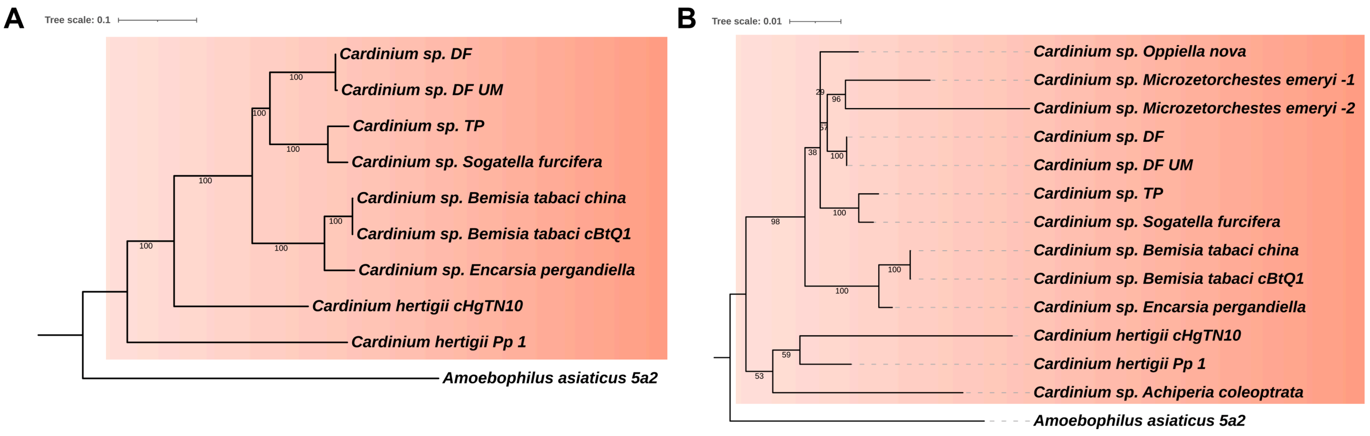

**Fig. S2. Phylogenetic analysis of *Cardinium* species**

**(A)** The phylogenetic tree was constructed based on 295 single-copy orthogroups. The annotated protein sequences of species were assigned into orthogroups by OrthoFinder v2.5.4. **(B)** The phylogenetic tree was constructed based on 16S rRNA sequences. All the source of those sequences were listed in Table S2. Two phylogenetic trees were constructed by the program RAxML and edited by the online tool iTOL. All *Cardinium* sequences were highlighted in orange background.

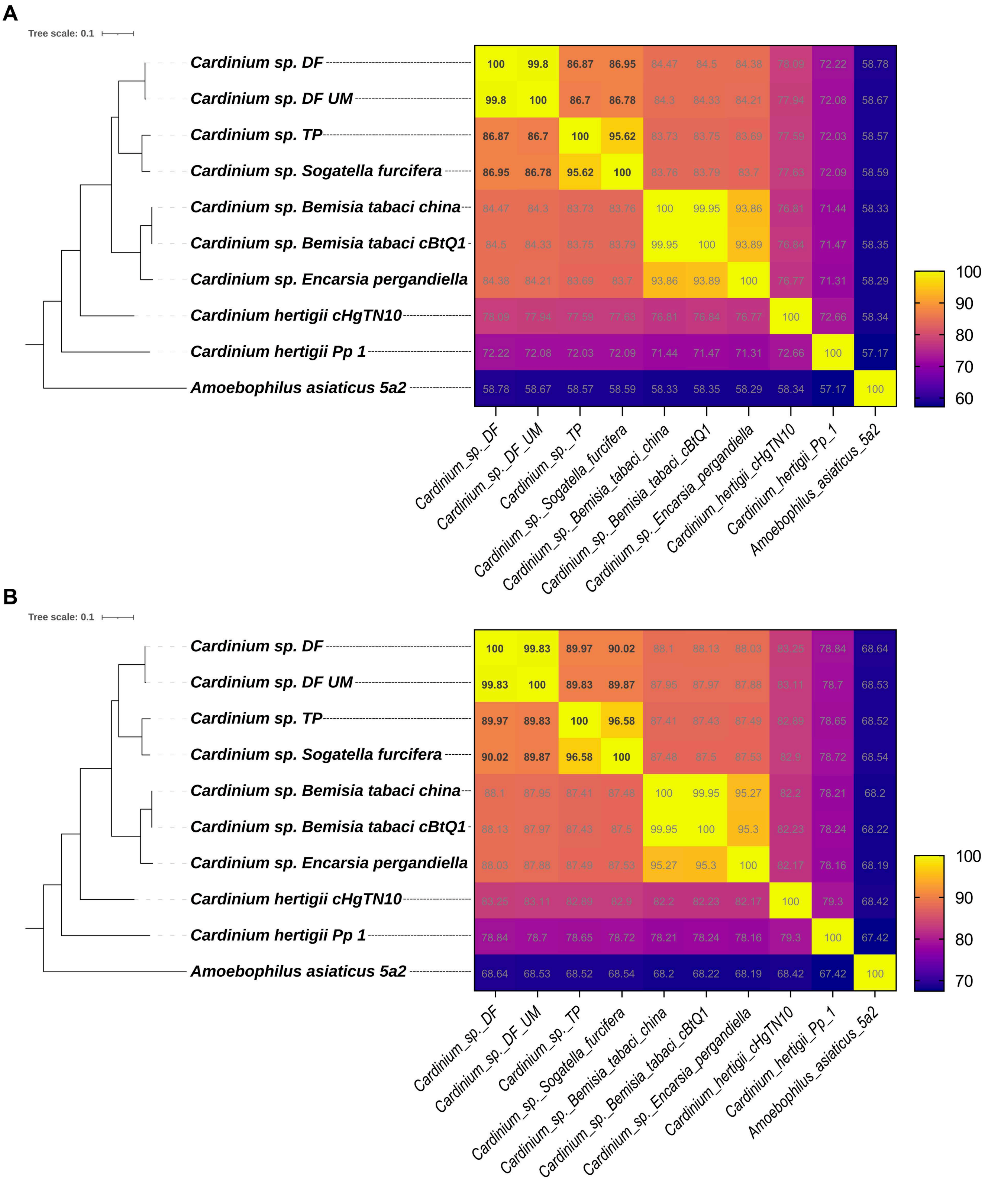

**Fig. S3. Identity and similarity matrices of *Cardinium* genomes**

The identity **(A)** and similarity **(B)** matrices of *Cardinium* genomes were constructed by the online tool SIAS (Sequence Identity And Similarity) in default parameters and based on the protein sequence alignment of genes in the 295 single-copy orthogroups assigned by OrthoFinder. The phylogenetic tree was adapted from Fig. S2A.

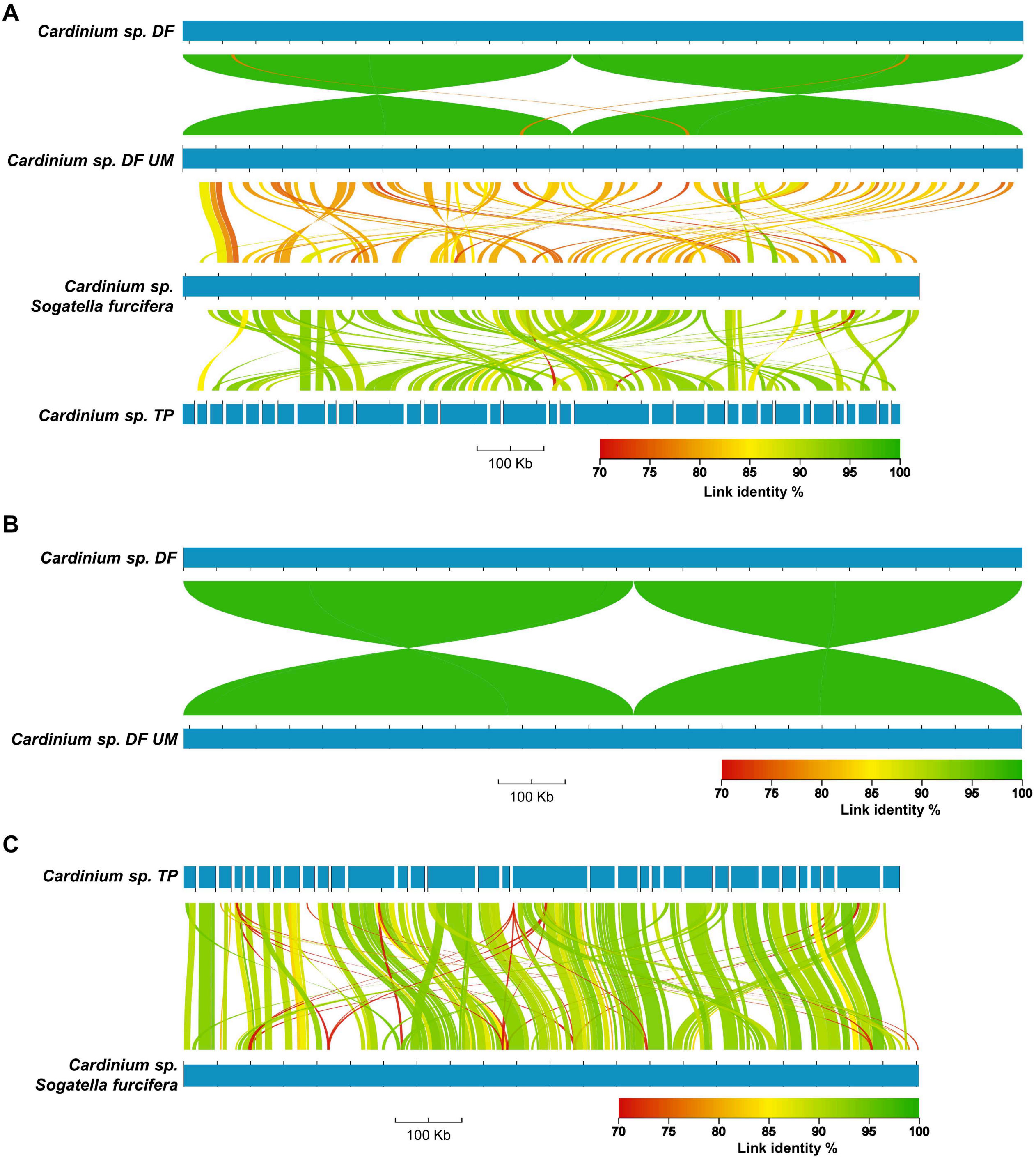

**Fig. S4. Whole-genome alignments of *Cardinium* genomes**

The whole-genome alignment of genome sequences were performed and presented by AliTV. **(A)** The whole genome alignment of four *Cardinium* species (Table 1 and S2) was filtered by 70% identity and 5-kb link length. Only the longest contig NZ_VMBH01000001.1 of *Cardinium sp. DF UM* was used in this alignment. **(B)** The genome sequences of two *Cardinium* sp. *DF* were aligned, and the alignment was filtered by 70% identity and 10-kb link length. **(C)** The genome sequences of *Cardinium sp. TP* and *Cardinium sp. Sogatella furcifera* were aligned and the alignment was filtered by 70% identity and 2-kb link length.

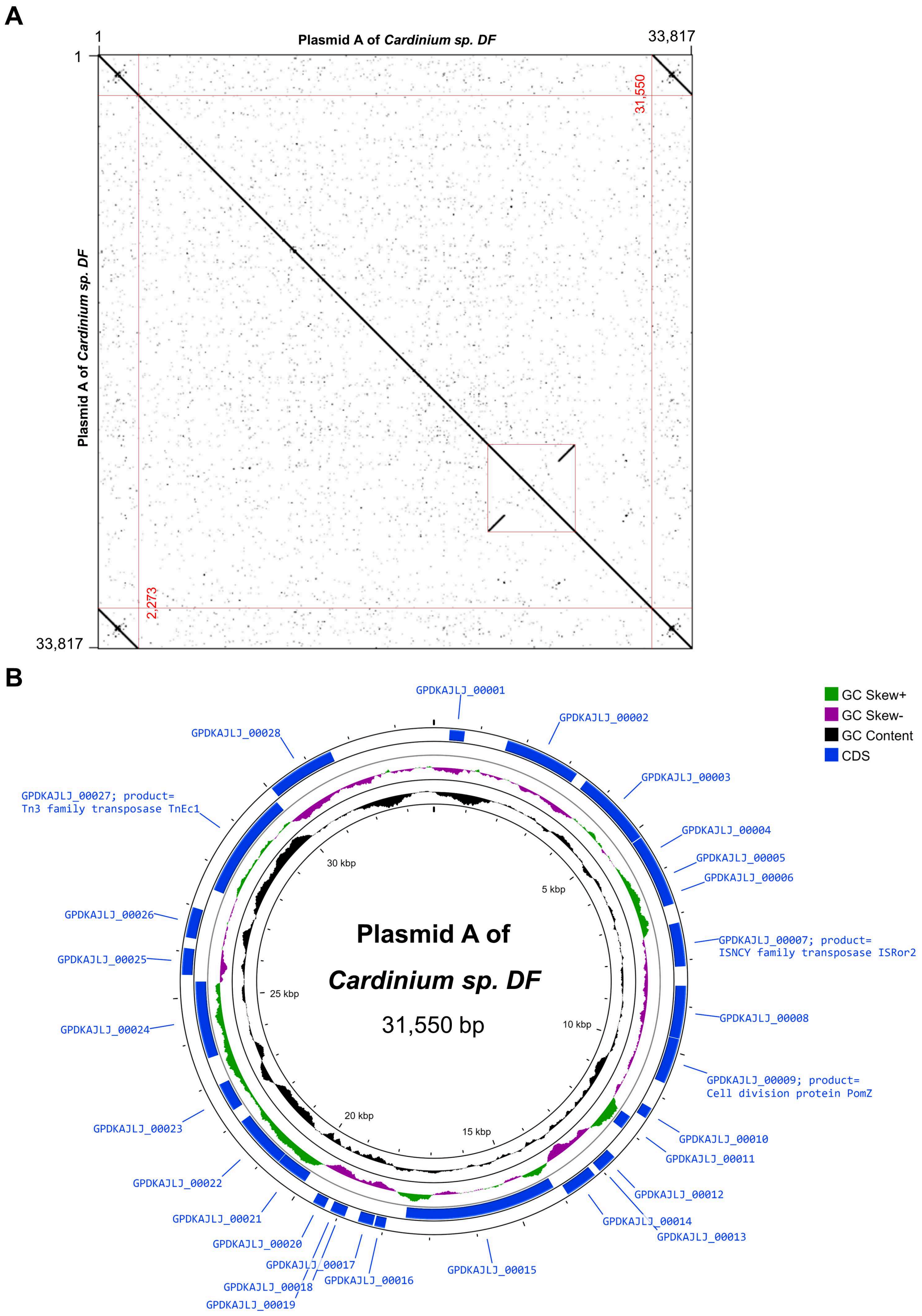

**Fig. S5. Dot plot and circular genome map of Plasmid A of *Cardinium sp. DF***
**(A)** Dot plot of the contig A (Fig. 1B) suggests the region 1-31,550 bp is a circular plasmid. This plasmid was named as Plasmid A of *Cardinium sp. DF* (Fig. 1A). **(B)** Circular genome map of Plasmid A of *Cardinium sp. DF*. The gene annotation was performed by Prokka and visualized by the online tool Proksee.

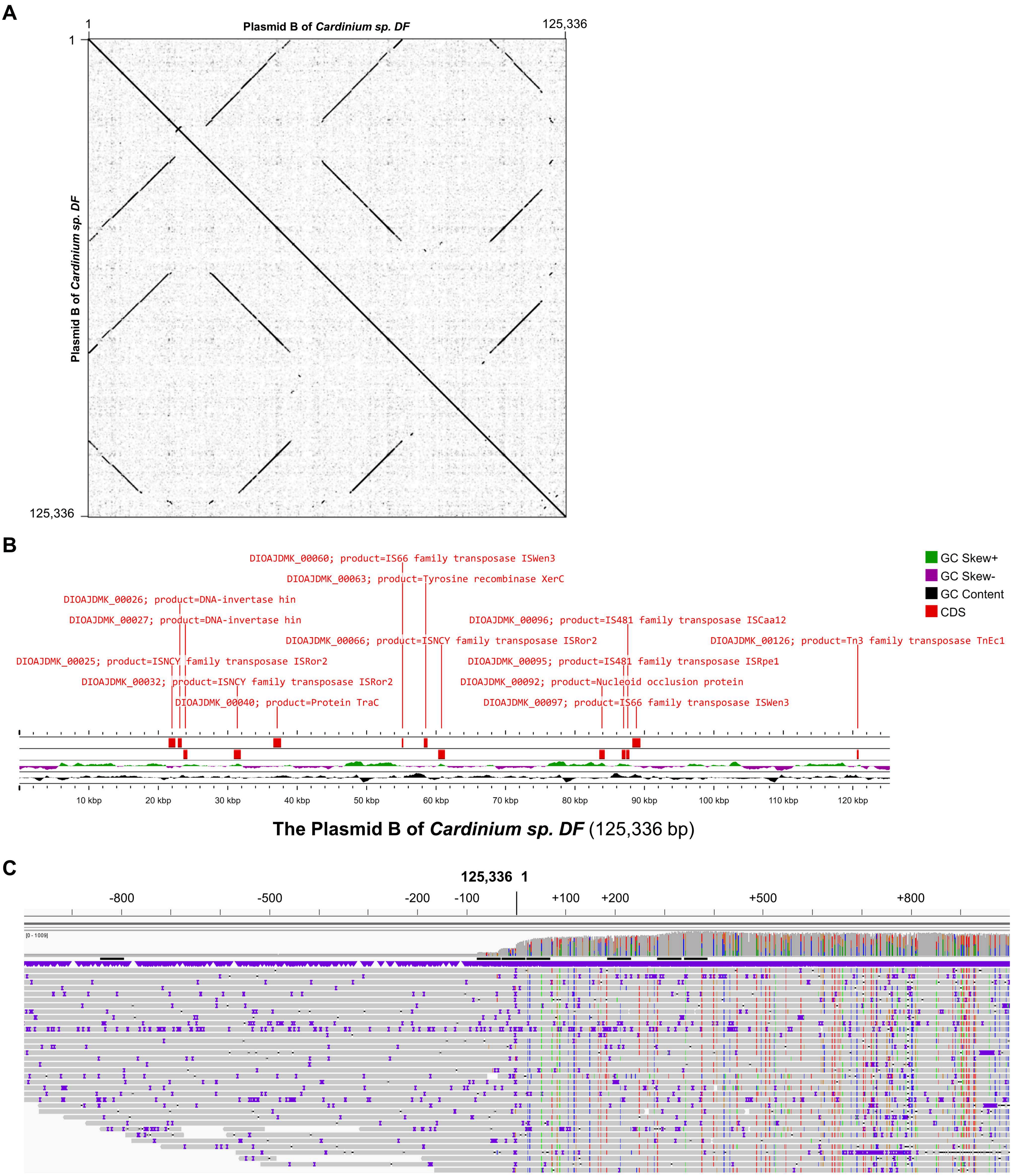

**Fig. S6. Dot plot and annotation of Plasmid B of *Cardinium sp. DF***

**(A)** Dot plot of Plasmid B of *Cardinium sp. DF* revealed a wide range of repeated sequences. **(B)** Linear genome map of Plasmid B of *Cardinium sp. DF*. The gene annotation was performed by Prokka and only those genes with functional names were visualized by the online tool Proksee. **(C)** TGS long reads mapping of the junction site including the 2.5 kb sequences on two ends. Over 80 long reads spanned the junction site and suggested this is circular plasmid DNA. Reads mapping was performed by Integrative Genomics Viewer (IGV).

**A**

```
DIOAJDMK_00108      -------MDNNQIKRKILFAMFRLCILFY-LCLVVCSYMEHRTGMLPNKNKIPIQEQSEC      52
GPDKAJLJ_00014      -------------------MFRLCILFY-LCPVACSYMAHRTGMLPNKNPMPSQVQLEY      39
LFAMDCML_00833      MRIIYYIMDNNQIKRKILFAMFRLCILFYLLCLVSCSYMAHRPGMLPNKNPIPIQEQSEC      60
                                       ********* ** * **** ** ******* :* * * *

DIOAJDMK_00108      GSTVCSYDSSTILKKRLKNKIKASVGVLLCYTNDKQESFVLLGRERVGKCDGGTWYELGG      112
GPDKAJLJ_00014      GSTVCSYDSKTILERRLEQNIKASVGVLLCYTNDKQESFVLLGRERVGKCDGETWCELGG      99
LFAMDCML_00833      GSTVCSYDSKTILERRLKNNLKASVGVLLCYTNDKQENFLLLGRERVGKCDGETWCELGG      120
                    *********.***::**::::****************.*:************ ** ****

DIOAJDMK_00108      SVETGESFLEAAIRESKEESGGIYQLDPNDLLNNGVVCYSEHKKKDGCIREEVYILLRVN      172
GPDKAJLJ_00014      SVETGESFLEAAIRESKEESEGIYQLDPNDLLNNGVVCYSEHKKKDGCIREEVYILLKVN      159
LFAMDCML_00833      SVETGESFLEAAIRESKEESGGVYQLDANNLLNNGVVCYNEHKKKDGCIREEVYILLKVN      180
                    ******************** *:**** *:*********.*****************:**

DIOAJDMK_00108      DFYPADVLCKAVNQQDKDTYREKDNFKWISCDSLCKIEHNPCVLKDIDGNEYTFMLRTFF      232
GPDKAJLJ_00014      DFYPADVLCKAVNHQNKDTYREKDDFKWVSCDSLCKVEHNLCVLKDIDGNAYTFTIRKFF      219
LFAMDCML_00833      DFYPADILCKAVNQQNKDTYREKDDFKWVSCDSLCTVEHNPCVLKDIDGNEYTFTLRKFF      240
                    ******:******:*:********:***:******.:*** ********* *** :*.**

DIOAJDMK_00108      YDALKRDSFLVKLKPVM   249
GPDKAJLJ_00014      YDALKRDAFLVKLKSVM   236
LFAMDCML_00833      YDALKRDSFLAKLKSVM   257
                    *******:**.*** **
```

**B**

```
AIMNMLDK_00972      MSSNNPFQNINQKQASISLQEHCEQAIVRKKIGRPLRPNMGKYLIKMDKGLHAKLSKYAE      60
DIOAJDMK_00064      MSSNNPFQNINQKQASISLQEHCEQAIVRKKIGRPIRPNMGKYLIKMDKGLHTKLSKYAE      60
                    ***********************************:****************:*******

AIMNMLDK_00972      ELGINNSFIISKALKEFFSKHCI      83
DIOAJDMK_00064      ELGISNSFIISKALKEFFSKYCI      83
                    ****.***************:**
```

**C**

```
AIMNMLDK_00971      MQILVANIKGGCGKSTLVSALADLLDADIIDHDNQGTIRVSSSFTGLRVPVTYEKITKKI      60
DIOAJDMK_00065      MQILVANIKGGCGKSTLVSALADLLDADIIDHDNQGTIRVSSSFTGLRVPVTYEKITKKI      60
                    ************************************************************

AIMNMLDK_00971      VLHDTPPYNTGNLKSLIQEADLILIPCKLMYPDLLALRALSDKLKKLNMTQKGVIVFNEI      120
DIOAJDMK_00065      VLHDTPPYNTGNLKSLIQEVDLILIPCKLMYPDLLALRALSDELKKLNMTQKAVIVFNEI      120
                    *******************.**********************:*********.*******

AIMNMLDK_00971      RKPHNNTYKEVKALYESNYSEIKKAKTELSNLISFSRVLSEPITGKALEQIKSLISELNI      180
DIOAJDMK_00065      RKPHNNTYKEVKALYESNYSEIKKAKTELSNLISFSRVLSEPVTGKALEQVKSLISELSI      180
                    ******************************************:*******:*******.*

AIMNMLDK_00971      Y   181
DIOAJDMK_00065      Y   181
                    *
```

**Fig. S7. Sequence alignment of reductive genes in *Cardinium sp. DF***

**(A)** Sequence alignment of the NUDIX hydrolase genes, LFAMDCML_00833 of *Cardinium sp. TP*, GPDKAJLJ_00014 of Plasmid A and DIOAJDMK_00108 of Plasmid B. **(B)** Sequence alignment of the hypothetical protein genes, AIMNMLDK_00972 of *Cardinium sp. Sogatella furcifera* and DIOAJDMK_00064 of Plasmid B. **(C)** Sequence alignment of NUBPL iron-transfer P-loop NTPase genes, AIMNMLDK_00971 of *Cardinium sp. Sogatella furcifera* and DIOAJDMK_00065 of Plasmid B. All sequence alignments were performed by the online tool Clustal Omega.

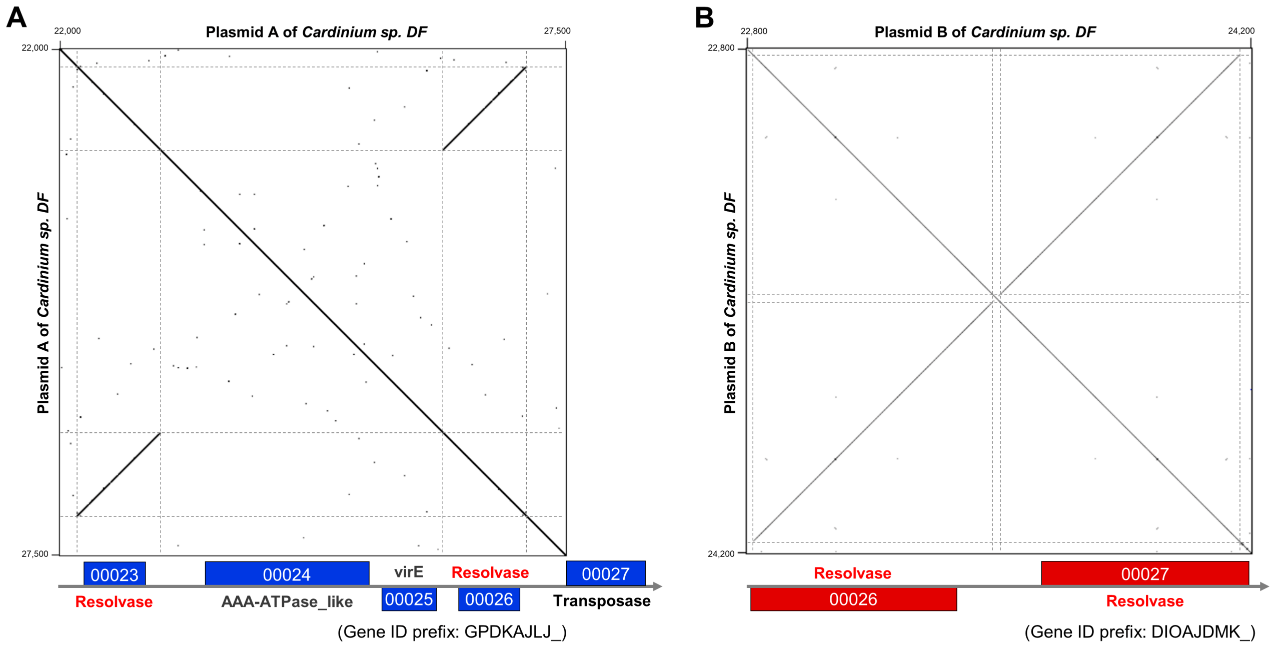

**Fig. S8. Dot plots of the inverted repeated regions in the two plasmids of *Cardinium sp. DF***

**(A)** Dot plot of the region 22,000-27,500 bp of Plasmid A of *Cardinium sp. DF*. The gene annotation was adapted from Table S3. **(B)** Dot plot of the region 22,800-24,200 bp of Plasmid B of *Cardinium sp. DF*. The gene annotation was adapted from Table S6. Two pairs of inverted resolvase genes were highlighted in red color.

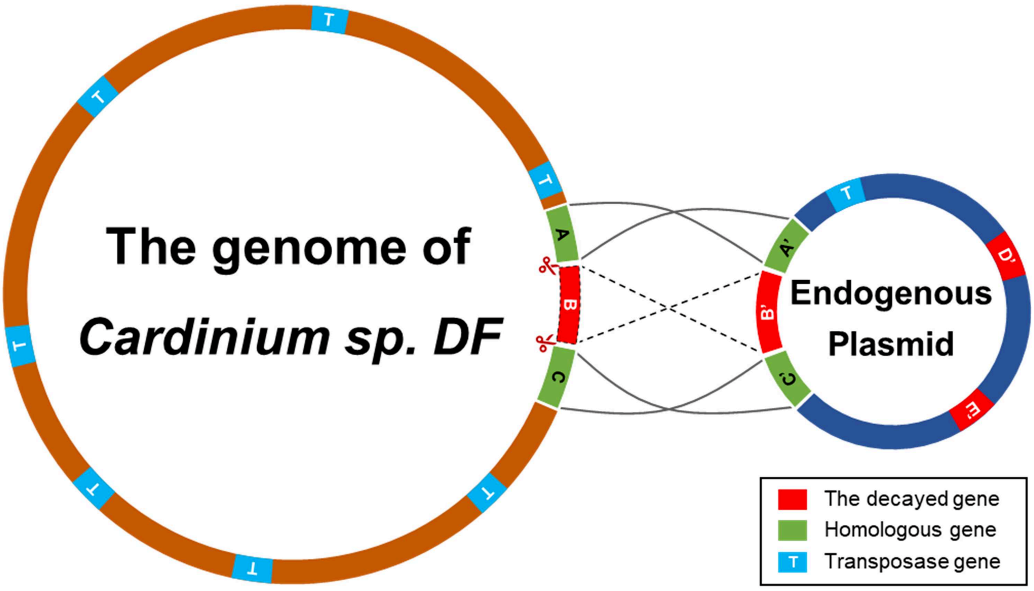

**Fig. S9. Graphical illustration of the genome reduction driven by endogenous plasmid**

In the absence of chromosomal recombination, the chromosomal genome of *Cardinium sp. DF* shared conserved or homologous genes and regions with the endogenous plasmid. When genes decay in the chromosomal genome (like the gene B), the homologous gene (the gene B') in plasmid could maintain the necessary cellular function and thus underpin the gene decay. Massive copies of transposase genes may participate in the molecular process of the genome reduction with endogenous plasmid.
